# Evidence for the Scarr-Rowe effect on genetic expressivity in a large US sample

**DOI:** 10.1101/429860

**Authors:** Michael A. Woodley of Menie, Jonatan Pallesen, Matthew A. Sarraf

## Abstract

Using the Continuous Parameter Estimation Method (CPEM), a large genotyped sample of the population of Wisconsin, USA (the Wisconsin Longitudinal Study, *N*=8,509) is examined for evidence of the Scarr-Rowe effect, an adverse gene x environment (GxE) interaction that reduces the heritability of IQ among those with low socioeconomic status. This method allows the differential expressivity of polygenic scores predictive of both educational attainment and IQ (EA3) on the phenotype of IQ to be operationalized throughout the full range of these variables. Utilizing a parental SES factor-weighted composite as a measure of childhood SES, evidence for the Scarr-Rowe effect was found, i.e. the genetic expressivity of EA3 on IQ increased with increasing parental SES (*β*=.064, *p*=3.82×10^−9^, *df*=8508). The effect was found for both the male and female samples separately (*β*(males)=.051, *p*=.002, *df*=4062; *β*(females)=.076, *p*=1.76×10^−7^, *df*=4445) — there were no significant sex differences in the effect magnitudes, however. The effects were furthermore robust to removing outlying values of parental SES.

## Introduction

The Scarr-Rowe effect is the apparent tendency of the heritability of IQ to be lower among those with lower compared to those with higher socioeconomic status (SES). This effect possibly results from adverse bioecological conditions restricting the variance in developmental opportunities among those with low SES, which, in turn, supresses the expression of IQ-related genetic variants via a gene x environment (GxE) interaction (Bronfenbrenner & Ceci, 1994). The effect was first described by Sandra Scarr (1971) in a study of Philadelphia school children (Scarr-Salapatek, 1971), and was subsequently replicated in the US population by David Rowe and colleagues (Rowe et al., 1999). Eric Turkheimer and colleagues reported one of the largest Scarr-Rowe effects, finding that among those with the lowest SES, the heritability of IQ was close to zero (Turkheimer et al., 2003).

A meta-analysis of 43 effect sizes sourced from 14 Scarr-Rowe effect studies found clear indications of a geographic clustering of the effects (Tucker-Drob & Bates, 2015). A significant Scarr-Rowe effect (operationalized as a genetic additivity x SES interaction) was present among the US population (*ρ*=.074, *SE*=.025, *p*=.003); but among populations sampled from Western Europe and Australia, the effect was absent. Even removing the US effect sizes reported by Turkheimer’s group (which were among the largest observed) yielded a significant Scarr-Rowe effect for this country (*ρ*=.058, *SE*=.020, *p*=.003), and also significant differences between the US and non-US samples (*Δ*=.085, SE=.030, *p*=.005). This meta-analysis also yielded clear indications of rising heritability with age, with the Scarr-Rowe effects being smaller among samples in which IQ had been measured at a later age (reflecting increasing influence from additive genetic effects). A recently published large *N* (1,636,968) genetically informed study of the population of Florida, containing 24,640 twins and 274,786 siblings, found no evidence for the Scarr-Rowe effect, however, even though the sample used was highly socioeconomically heterogeneous and consisted of children (Figlio et al., 2017).

The recent availability of high-quality polygenic scores (PGS; normally distributed genetic indices comprised of variants that collectively significantly predict variance in a trait of interest) for educational attainment and also IQ enables a new method for estimating GxE interaction effects, as their expressivity with respect to these phenotypes (i.e. the degree to which the genotype influences the phenotype) can be estimated, yielding more direct indications of such interactions that overcome certain limitations inherent in the twin design (such as ambiguities regarding the causation of phenotypic convergence and divergence; see e.g. Segal, 2013). So far one study has already employed genomic data in investigating the Scarr-Rowe effect (Tahmasbi et al., 2017). In this study, a maximum likelihood model was developed to examine the presence of Scarr-Rowe effects in a sample of 40,172 individuals — sourced from the UK BioBank database for whom Genome-wide complex trait analysis Genome-based restricted maximum likelihood estimates of heritability were computed for individuals of different SES – where it was found that genetic variance in IQ increased as SES decreased, yielding an anti-Scarr-Rowe effect, which, as the authors note, is consistent with the null and negative effects typically reported outside of the US (Tucker-Drob & Bates, 2015).

In the present study, a large and socioeconomically representative genotyped sample of the state of Wisconsin (the Wisconsin Longitudinal Study; WLS) will be used to investigate the presence of Scarr-Rowe effects via the application of a novel (and very straightforward) method that permits the direct operationalization of the expressivity of PGS on IQ, which will be used to determine the presence of GxE interactions as a function of parental SES.

## Methods

### Sample and measures

All data were collected from the WLS, a longitudinal study of a randomly selected sample of Wisconsin High School students born between 1937 and 1940, which began data collection in 1957 (when the participants were in their 30s); the most recently collected data wave is from 2011. The sample is nearly exclusively of European descent, consistent with its high representativeness of mid-century Wisconsin demographics (Herd et al., 2014).

### Polygenic score for general intelligence

In the period 2007/8 and again in 2010, a large genetic data collection exercise was undertaken in which saliva samples were obtained from a total of 9,012 individuals, who were subsequently genotyped using the Illumina *HumanOmniExpress* array as part of a very large genome wide association study, examining variants predictive of individual differences in educational attainment and related cognitive phenotypes (Lee et al., 2018). For full information on genotyping procedures, see: https://www.ssc.wisc.edu/wlsresearch/documentation/GWAS/Herd_QC_report.pdf. Several alternative PGS were released, each representing different collections of phenotypes against which variants had been regressed, and also different methods (e.g. GWAS vs. MTAG – a multivariate regression-based estimation method). One PGS was selected for the present analysis, PGS_EA3_MTAG (henceforth EA3), which was trained via multivariate analysis with respect to several convergent cognitive phenotypes, including an IQ test from UK BioBank, various neuropsychological functioning tests and IQ subscales from COGENT, self-reported mathematical ability, and highest mathematics class successfully completed. Finally, also included among the training phenotypes was educational attainment, which was defined based on the 1997 ISCED UNEASCO classification, which ranks individuals based on seven internationally comparable categories of educational attainment, rescaled in terms of US equivalent years-of-schooling. EA3 comes closest to capturing variance with respect to an overarching general intelligence factor. It should finally be noted that the sample from which Lee et al. (2018) derived their PGS was extremely large (*N*>1 million; WLS was but a small part of the overall sample) and was ethnically heterogeneous, thus the PGS were corrected for population stratification. This coupled with the extremely high ethnic homogeneity of the WLS sample (Herd et al., 2014) eliminates the need to include additional controls for population stratification in analyses utilizing these scores.

### Henmon-Nelson test of mental ability

The WLS contains participant scores on the Henmon-Nelson IQ test (variable code: GWIIQ_BM), which measures the domains of spatial, verbal and mathematical ability. The test is timed, taking 30 minutes to complete, and consists of 90 items presented in ascending order of difficulty. The test was standardized state-wide in Wisconsin during the initial 1957 data collection wave, when the participants were in their 30s. The test exhibits excellent psychometric characteristics, including high internal consistency (α≈.95; Hansen, 1968; Harley, 1977) and also high convergent validity with respect to other measures of IQ, correlating in the *r*≈.80 - .85 range with Fullscale IQ as measured using the WAIS (Klett & Watson, 1986; Kling et al., 1978).

### Parental SES

The WLS contains a factor-weighted composite measure of parental SES (SES57). This measure is comprised of father’s years of schooling (EDFA57), mother’s years of schooling (EDMO57), Duncan’s socioeconomic index for father’s 1957 occupation (OCSF57), and average parental income, with estimates for missing data (PI5760). All data were collected in the 1957 wave.

### Sex

Data on the sex of WLS respondents (SEXRP) were collected in order to test for sex differences in the magnitudes of any Scarr-Rowe effects that might be present. This variable was measured in 1957, with 1 = male and 2 = female.

### Analytical strategy

#### Continuous Parameter Estimation Model

For the present analysis the Continuous Parameter Estimation Method (CPEM) will be used to test for Scarr-Rowe effects on genetic expressivity. CPEM was developed by Richard Gorsuch (2005) and is based on the mathematics of the Pearson Product Moment correlation. The formula for the Pearson correlation can be written as *r*=Σ(*zx*zy*)/*N*, where *z*x and *z*y are the standardized scores for the independent and dependent variables respectively, *z*x∗*z*y is the dot product term for the two – the average of which across subjects yields the correlation coefficient *r*. Gorsuch (2005) proposed that the product term (*zx*zy*) for each individual is mathematically equivalent to a correlation for an *N* of 1. Thus, the dot product term functions as a continuous parameter estimate (CPE) of the covariance between the independent and dependent variable and can be used in regression models along with other variables for moderation analysis.

A very fruitful application of this technique has been in examining ability differentiation, such as the cognitive and strategic (behavioral) differentiation-integration effort effects, where covariance among clusters of cognitive abilities or behavioral indicators is expected to vary as a function of participants’ life history speed (Figueredo et al., 2013; Woodley et al., 2013). CPEM has also been utilized in a variety of other contexts, including the estimation of individual-level heritabilities derived using the correlational Falconer’s formula, for the purpose of examining whether the heritability of the latent life history *K* factor decreases as level of *K* increases (Woodley of Menie et al., 2015). CPEM has been used as an alternative to the method of correlated vectors in establishing latent variable moderation effects (Woodley of Menie et al., 2015), in quantifying the impact of age on assortative mating on emotional intelligence (Śmieja & Stolarski, 2018), in examining the role of SES as a moderator of the association between stable life history strategy and sexual debut (Dunkel et al., 2015), and for examining the curvilinear associations between longitudinal trends among the WAIS scale-scores and participant age (Lee et al., 2008), among other things.

Here it is proposed that CPEM can be used to compute the individual-level covariance among EA3 PGS and IQ scores for each participant utilizing the dot product terms to capture the strength of the association between subject genotype and phenotype, which is a direct measure of genetic expressivity. By regressing parental SES against the CPE, the presence of a Scarr-Rowe effect can be determined if the resulting *β* value is positive, as this would indicate that the genetic expressivity (i.e. the covariance) of EA3 PGS to IQ increases as parental SES increases. Given that large amounts of data are available in WLS for both sexes, comparison of the effect sizes can be used to determine the presence of a sex difference in the effect. All analyses are conducted in R and the code is publicly archived at: http://rpubs.com/Jonatan/cpem

## Results

### Analysis 1: Combined sample

Table 1 presents the descriptive statistics and correlations for the variables utilized in the analysis of the combined sample.

**Table 1.** Descriptive statistics and correlations for the combined sex sample.

| **Descriptives** | *PGS* | *IQ* | *Parental SES* |
| --- | --- | --- | --- |
| *N* | 8,509 | 10,317 | 10,317 |
| *Min.* | -.891 | 61 | 1 |
| *Max.* | .351 | 145 | 97 |
| *Mean* | -.285 | 100.5 | 15.98 |
| *Std. Dev.* | .154 | 14.92 | 11.03 |
| **Correlations** |  |  |  |
| *PGS* | 1 |  |  |
| *IQ* | .26 | 1 |  |
| *Parental SES* | .15 | .30 | 1 |

*Note: All correlations significant at the <.0001 level*

More information on the descriptives is available here: http://rpubs.com/Jonatan/cpem. Table 2 presents the results of the CPEM analysis. As is standard in analyses involving CPEM, all variables are standardized prior to entry into the regression (e.g. Figueredo et al., 2013). This means that the resultant *b* values correspond to standardized *β* values, and no intercept term needs to be computed, yielding one additional model degree of freedom. In addition to the CPEM regression parameter, the model residual skewness is also estimated in order to ensure that there are no normality violations.

**Table 2.** The results of regressing the CPE z(*EA3*)\**z*(IQ), which captures differences in levels of genetic expressivity on parental SES. The model *t*-statistic, significance, degrees of freedom and the skew on the residual of parental SES are also presented. *Adj*. *R*^*2*^=.004, *F*=34.79.

| Parental SES | $\beta$ (SE) | $t$ | $p$ $r(>\vert t\vert )$ | $Df$ | $z$ (residual SES) |
| --- | --- | --- | --- | --- | --- |
| $z(EA3)*z(IQ)$ | .064 (.011) | 5.90 | $3.82 \times 10^{-9}$ | 8508 | 1.14 |

The regression model yields indications of a small-magnitude (i.e. <.29; Cohen, 1988) Scarr-Rowe effect, when parental SES is used to predict variation in the genetic expressivity of the participant’s PGS on their IQ scores. The effect is highly statistically significant (which is unremarkable given the very high model degrees of freedom) and the skew on the model residual falls within the levels generally considered acceptable for parametric regression (i.e. *z* between +2 and −2; George & Mallery, 2010). The results of this analysis are graphed in Figure 1.

**Figure 1.**
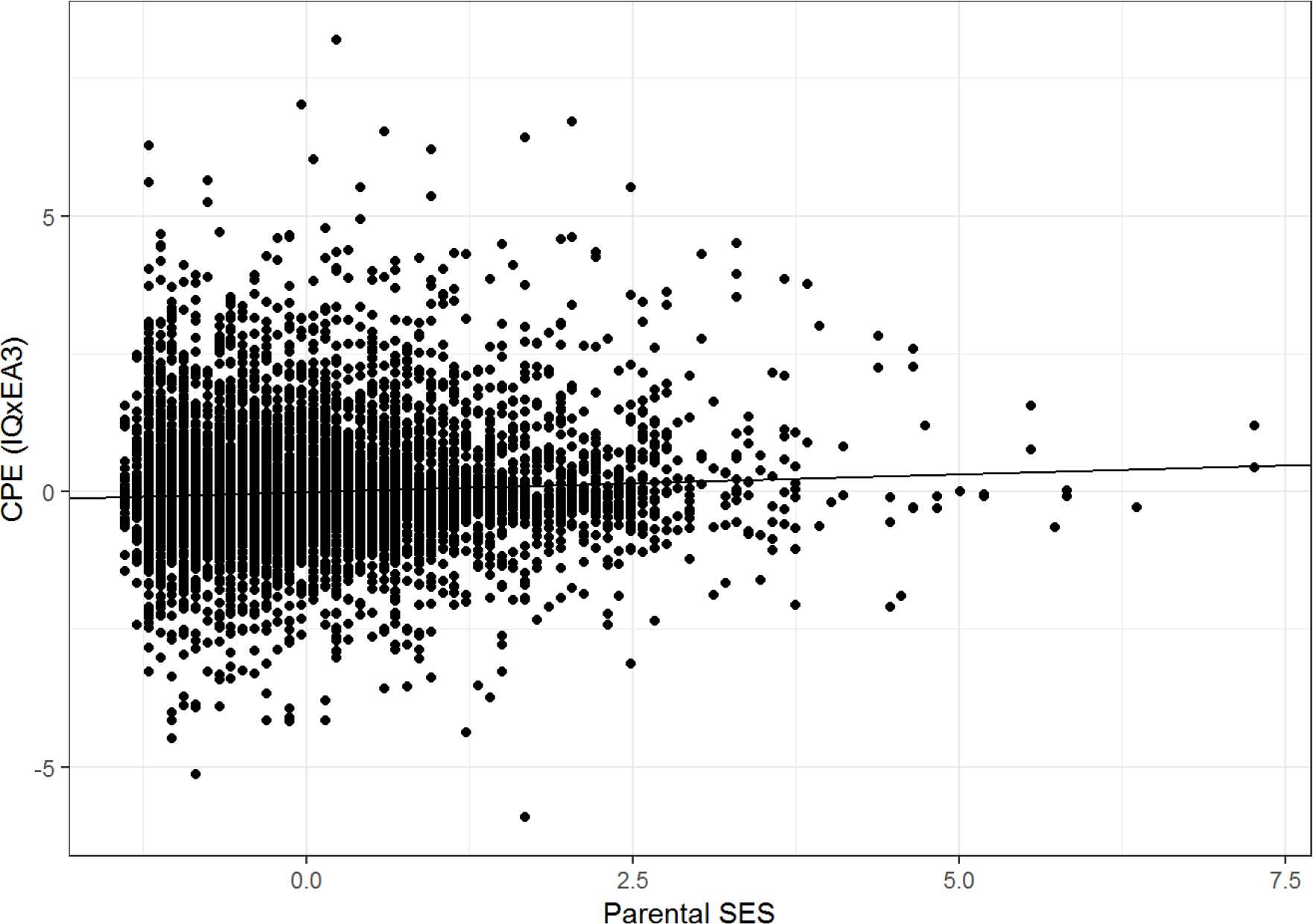
Scatter plot and regression line of the CPE z(*EA3*)\**z*(IQ) capturing individual differences in genetic expressivity as a function of parental SES for the combined sample, *N*=8,509.

### Analysis 2: Broken out by sex

Table 3 presents the correlations broken out by sex.

**Table 3.** Correlations broken out by sex, with males below the diagonal and females above.

|  | *PGS* | *IQ* | *Parental SES* |
| --- | --- | --- | --- |
| *PGS* | 1 | .27 | .16 |
| *IQ* | .26 | 1 | .29 |
| *Parental SES* | .14 | .31 | 1 |

*Note: All correlations are significant at $p < .0001$ .*

*Note: All correlations are significant at p<.0001*.

Table 4 presents the results of CPEM analysis for males and females separately. In addition to the CPEM regression parameter, the model residual skewness is also estimated for both regressions in order to ensure that there are no normality violations.

**Table 4.**
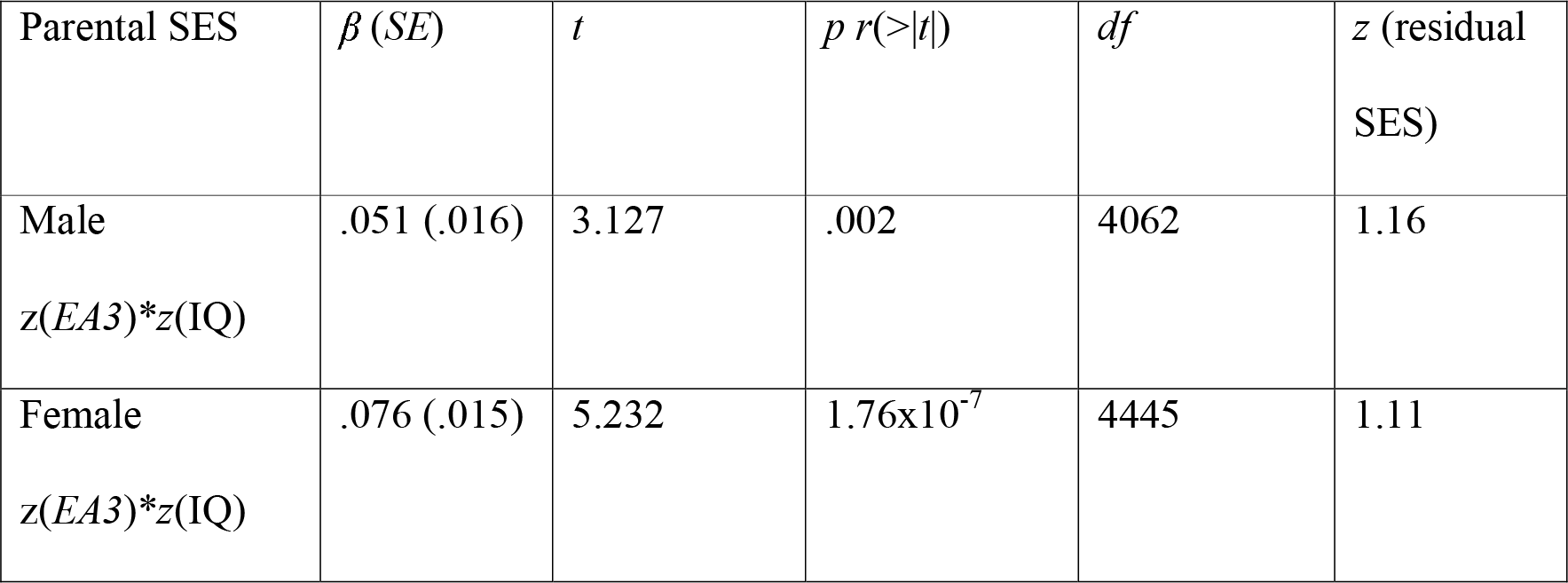
The results of regressing the CPE z(*EA3*)\**z*(IQ), which captures differences in levels of genetic expressivity against parental SES for males (top row) and females (bottom row) separately. The model *t*-statistics, significances, degrees of freedom, and the skew on the residuals of parental SES are also presented. Males *Adj. R^2^*=.002, *F*=9.776. Females *Adj. R^2^*=.006, *F*=27.37.

The effect is present in both males and females separately and, as with the combined sample, the residual model skewness falls within the acceptable range of values (i.e. *z* between +2 and −2). The female *β* is larger in magnitude than the male *β*; however, the difference in magnitude is not statistically significant (*z*=.123, *p*=.246), indicating no sex differences in the effect sizes.

### Robustness analysis: Outlier removal

To test the robustness of the effects to potentially outlying values of parental SES the analyses were rerun for the combined, male and female subsamples respectively excluding all values of parental SES that were ≥+4SDs above the mean. The results of this analysis are presented in Table 5.

**Table 5.** The results of regressing the CPE z(*EA3*)\**z*(IQ), which captures differences in levels of genetic expressivity against parental SES for the combined sample (top row), males (middle row) and females (bottom row) separately. The model *t*-statistics, significances, degrees of freedom, and the skew on the residuals of parental SES are also presented. Combined sample *Adj. R^2^*=.003, *F*=23.15, Males *Adj. R^2^*=.002, *F*=7.62. Females *Adj. R^2^*=.003, *F*=16.43.

| Parental SES | $\beta$ (SE) | $t$ | $p$ $r(>\vert t\vert )$ | $Df$ | $z$ (residual SES) |
| --- | --- | --- | --- | --- | --- |
| $z(EA3)*z(IQ)$ | .050 (.010) | 4.81 | $1.52 \times 10^{-6}$ | 8423 | .748 |
| Male<br>$z(EA3)*z(IQ)$ | .042 (.015) | 2.76 | .006 | 4021 | .736 |
| Female<br>$z(EA3)*z(IQ)$ | .057 (.014) | 4.05 | $5.12 \times 10^{-5}$ | 4401 | .762 |

A more stringent outlier cut-off was also employed in a second outlier removal analysis (≥+3 SDs above the mean). This had no influence on the effect direction, although in the case of the male sample, the *β* was no longer statistically significant. The results of this additional analysis can be viewed here: http://rpubs.com/Jonatan/cpem. The results indicate that outlying values of parental SES are not driving these effects.

## Discussion

An analysis using CPEM indicates the presence of an apparent Scarr-Rowe effect on the genetic expressivity of a PGS capturing variance in general intelligence on a phenotypic measure of intelligence, in a well-powered study. Genetic expressivity increases as SES increases, consistent with the findings of gene x SES interaction effects from US cohorts (Tucker-Drob & Bates, 2015), which means that the present method of estimating the effect using the differential expressivity of PGS on IQ as a function of SES yields equivalent results to studies estimating gene x SES interaction effects derived using more conventional behaviour genetic approaches (i.e. Biometric Structural Equation Modelling using twins and siblings). Furthermore, to the best of our knowledge, this is the first time that the possibility of sex differences in the Scarr-Rowe effect has been investigated; but even though the effect is larger in females than in males, the difference was not statistically significant.

While these findings are supportive of the existence of the Scarr-Rowe effect and are broadly congruent with relevant studies from the US, there is evidence that not all parts of the US are equally conducive to the effect, possibly due to high SES variation both within and among US States. The Florida cohort study of Figlio et al. (2017) is illustrative on this score, as it was extremely highly powered to detect the effect yet found no indications of an SES x IQ heritability interaction using both twins and siblings. The most conservative interpretation of our results therefore is that the bioecological factors that supress the expressivity of cognitive genetic variants among those with low levels of childhood SES were present specifically among those born in the State of Wisconsin in the late 1930s and early 1940s. From this arises the question of whether there might be a secular trend in the strength of the Scarr-Rowe effect. Perhaps one reason that Figlio et al. (2017) were unable to detect the effect is that environmental quality in the US among those with low SES has improved in the decades since the WLS cohort was born (the Figlio et al., 2017 cohorts were born between 1994 and 2002, approximately six decades later), thus erasing the effect. A cross-temporal meta-analysis of the US data in Tucker-Drob and Bates (2015) along with Figlio et al. (2017) might help to determine whether such a trend exists, net of factors such as participant age at cognitive evaluation and location within the US. It would also be interesting to see if the strength of genetic expressivity utilizing the present methods increases when IQ is measured using younger cohorts, which would also be consistent with findings from Tucker-Drob and Bates (2015) of a “fading out” of the effect when heritability and IQ are evaluated in older cohorts.

Finally, given that *g* has a potentially very flat norm of reaction (meaning that the trait seems to be well canalized against environmental and certain genetic influences experienced during development and childhood; Protzko, 2017; Sesardic, 2005), it is predicted that the biggest and, critically, most persistent impact of bioecological elicitors of the Scarr-Rowe effect will be on measures of IQ exhibiting low *g* saturation, and thus low heritability (see, e.g., Voronin et al., 2015), which potentially leaves greater “room” for GxE interactions in the determination of trait variance. Where environmental deprivation might be expected to have its greatest impact is on those narrower and more trainable specialized manifestations of intelligence, which will be more reliant upon the presence of opportunities that may boost heritability via developmental opportunities present in the child’s environment that enhance genetic expressivity of variants underlying intelligence. If it is found that *g* loading negatively moderates ability measures’ sensitivity to the Scarr-Rowe effect, then the *g* loading of tests might be an important factor to control for in future meta-analyses. Moreover, it suggests that the Scarr-Rowe effect may help increase our understanding of the Flynn effect (which also occurs to the greatest extent on the least *g*-loaded abilities; te Nijenhuis & van der Flier, 2013), as reductions in the strength of the former effect may be a driver of the latter effect. This is because reduced variance in the provisioning of environmental factors such as educational attainment and other inducements towards cognitive specialization may be boosting opportunities for those with low SES to reach their genetic potential in terms of their capacity to cultivate specialized abilities, leading to potentially large gains in IQ, especially in instances where the transition from a poor- to a high-quality environment is very rapid. This may explain why in a substantial subset of studies, the Flynn effect appears to be larger among those with lower levels of IQ (which tracks lower SES) (e.g. Flynn, 2012).

